## supplementary information for "Modelling membrane reshaping by staged polymerization of ESCRT-III filaments"

October 3, 2022

[1] *Department of Physics and Astronomy, Institute for the Physics of Living Systems University College London, London, UK*
[2] *MRC Laboratory for Molecular Cell Biology, University College London, London WC1E 6BT, United Kingdom*
[3] *Institute of Science and Technology Austria, 3400 Klosterneuburg, Austria*
[4] *Department of Biochemistry, University of Geneva, Geneva, Switzerland*
[5] *MRC-Laboratory of Molecular Biology, Cambridge, United Kingdom*

#### Supporting Information

*

### S1 Electron microscopy

Protein purification is performed as described in Pfitzner et al. [1]. For LUV preparation, DOPC:DOPS:Rhodamine-PE (6:4:1; 10 mg/ml) mixture is evaporated in a glass tube, 500 $\mu$L of buffer are added, the tube is vortex followed by 5 times freezing and thawing. LUVs are then diluted 1:100 in buffer (20 mM Tris pH.6.8, 200 mM NaCl and 1 mM $MgCl_2$), spun down (10', 5,000 g), resuspended in 4.5 M Snf7 for 6h (4°C), before 1 $\mu$M Vps2, 1 $\mu$M Vps24, 2 $\mu$M Did2 and 2 $\mu$M Ist1 are added. Following overnight incubation at 4°C, all samples are spun down 10' 5,000 g and resuspended in buffer (negative stain EM) or buffer containing 30% glycerol (cryofreeze-fracture). Negative stain samples are absorbed onto EM grids and stained with 2% uranyl acetate for 30 s. Freeze fracture samples are transferred onto sample stamps, flash-frozen and processed using a 060 Freeze-Fracture System (BAF). Images are acquired on a Tecnai G2 Sphera (FEI) electron microscope.

### S2 Simulation protocols

#### S2.1 General settings

The three individual ESCRT-III filaments are each modelled using the 3-beaded model described in Harker-Kirschneck et al. [2]. The Flat Spiral has a target geometry of a flat ring which is bound to the membrane at an angle of 90° with respect to its axis of curvature ($\tau = 0°$ following the original notation in [2]). The Wide Helix and the Tight Helix have target geometries of tilted rings (i.e., no pitch is applied in the target geometries) which are bound to the membrane at an angle of 0° with respect to their axes of curvature ($\tau = 90°$ following the original notation in [2]). All neighbouring subunits within a filament are connected to each other via nine harmonic bonds with default bond constant $k = 256\ k_{\mathrm{B}}T/\sigma^2$, where $\sigma$ is the MD unit of length, unless stated otherwise. This determines the target geometry of the filament.

The initial structure of the membrane is a flat layer of membrane beads arranged in a hexagonal lattice. The membrane is modelled in its fluid state via the one-bead-thick model developed by Yuan et al. [3], with $\epsilon = 4.34\ k_{\mathrm{B}}T, \mu = 3.0, \xi = 4.0, \theta_0 = 0°$, which yields a fluid membrane of bending rigidity $\sim 20k_{\mathrm{B}}T$. Short-ranged 12-6 Lennard-Jones (LJ) interactions are applied between the two bottom beads of the filament's three-beaded subunits and the membrane beads. The membrane binding affinity of all the filaments is kept the same. Short-ranged LJ attractions are also applied at the interfaces of different filaments, through their neighbouring bottom beads (i.e. the inner bottom bead of the Flat Spiral with the outer bottom bead of the Wide Helix and the inner bottom bead of the Wide Helix with the outer bottom bead of the Tight Helix). The attraction is weak enough to allow detachment. The rest interactions within filament beads are volume exclusions.

For simulations with a generic cargo, we place the cargo particle in the centre of the filaments. A weak short-ranged LJ interaction is applied between the cargo particle and the membrane beads, with the interaction strength $\epsilon_{\mathrm{memb-cargo}} = 0.6\ k_{\mathrm{B}}T$. This allows the cargo to stay attached to the membrane, resisting fluctuations, but to not bud spontaneously in the absence of the filament. Volume exclusion is imposed between the cargo particle and the filament beads.

The radii of the coarse-grained beads used in our simulations are $r_{\mathrm{subunit}} = r_{\mathrm{membrane}} = 0.5\ \sigma, r_{\mathrm{cargo}} = 8.0\ \sigma$, where one subunit bead refers to one bead in the 3-beaded protein subunit. A conversion of $\sigma = 2.3$ nm is used to map the MD lengths to physical lengths [2]. All the short-ranged LJ interactions are cut-off at $r = 1.3\ r_{\mathrm{min}}$, where $r_{\mathrm{min}}$ is the distance at the potential minimum, and are shifted to zero at the cut-off. The default interaction strength $\epsilon = 3.0\ k_{\mathrm{B}}T$ is used to model all short-ranged LJ interactions for filament-membrane and filament-filament binding, unless specified otherwise. This interaction strength and cut-off range is strong enough to ensure filament-membrane association, but weak enough so that specific filament beads can interact with different membrane beads; thus filaments can slide over the membrane. All the volume-exclusion interactions are treated as LJ interactions truncated at the potential minimum and shifted to zero, with interaction strength $\epsilon = 2.0\ k_{\mathrm{B}}T$. All the interaction parameters are summarized in S1 Table.

The parameters in our model are the following: the radii of the three filaments, rigidities of the three filaments, the filament length, the strength of filament-membrane adhesion, and the diameter of the cargo particle. While we are not modelling a specific system, the geometrical parameters that we choose are in the regime of physiologically-relevant parameters. The target radii of the filaments (30 nm for Flat Spiral, 28 nm for Wide Helix) are based on the AFM measurements reporting Snf7 spirals to be 20-50 nm wide on membrane substrates [4, 5], and the cargo diameter (37 nm) is based on the reported ILV and exosomes diameters of 30-160 nm. The target radius of Tight Helix is chosen by a parameter scan, and it matches well with the theoretical fission limit ($\approx$ 3nm) [6]. The rigidities of the first two filaments are selected such that they yield the observed intermediate conical deformation [1] and, for simplicity, we use the same stiffness for the third filament. Little is known about the strength of the membrane adhesion of these composite filaments. The filament-membrane adhesion is chosen as the smallest value that allows the filaments to remain stably attached to the membrane. This parameter was previously tested when the single filament model was originally developed [2].

The simulations are run with molecular dynamics (MD) in the isothermal-isobaric ensemble with the barostat targeted at zero pressure. The barostat is applied in coupled x and y directions with damping factor of $10\tau$, where $\tau$ is the MD time unit. Langevin thermostat is applied at each MD step with the temperature set to 1 and the damping coefficient set to 1. Periodic boundary conditions are applied in the x and y directions. A time step of $0.01\tau$ is used. All MD simulations are carried out with LAMMPS code [7] and reduced units are used during the simulation. The unit conversion factors are summarized in S2 Table.

##### S2.2 Simulation of staged recruitment and disassembly of three filaments with cargo

We first equilibrate a generic cargo by itself on the membrane before adding the ESCRT-III filaments to the system. The membrane is constructed of 29440 beads ($160\times160\ \sigma^2$) and is equilibrated for $10^4$ steps, while the cargo and the filaments are kept frozen. Then we allow the cargo to move and equilibrate it for $10^6$ steps on the membrane, before placing the three ESCRT-III filaments around it in the shape of a three-stranded flat spiral. Each filament contains 84 subunits.

Although the copolymer is composed of three different filaments, at this stage, only the Flat Spiral is activated. Its target geometry is a flat ring with a target radius of 30.8 nm. The two inner filaments also behave like the Flat Spiral, thereby effectively remaining deactivated. The Wide Helix follows the same target geometry as the Flat Spiral, but with a slightly smaller target radius of 28.0 nm, since it is initially closely packed inside the Flat Spiral. This deactivation is further achieved, by temporarily turning the two inner filaments into one, by coupling their rigid body subunits (i.e., one subunit from the Wide Helix and one from the Tight Helix form a rigid body). This means the Tight Helix now behaves as if it were part of the Wide Helix (i.e., no target geometry for the Tight Helix), rather than an independent filament. We carry out the simulation for $10^6$ steps.

To simulate the first copolymerisation, we activate the Wide Helix by changing its target geometry from a flat to a tilted ring, that is 30.8 nm wide, while keeping the Tight Helix as part of it. After another $10^6$ steps, the Flat Spiral is disassembled instantaneously. This is modelled by deleting the harmonic bonds between protein subunits and resetting the membrane-filament interactions from short-ranged LJ interactions to volume exclusions.

After another $10^6$ steps, we simulate the shift in filament composition from the Wide Helix to the Tight Helix, by activating the Tight Helix to let it slowly constrict. For this we have deleted the rigid bonds between the Wide Helix and the Tight Helix and let them move independently. The Wide Helix remains in its previous state with a target radius of 30.8 nm. Meanwhile the Tight Helix starts off in exactly the same state as the Wide Helix, but progressively reduces its target radius from 30.8 nm to around 3 nm, which takes $2.5\times10^6$ steps in total. Half way through this stage, we instantaneously disassemble the Wide Helix, to prevent the Tight Helix from detaching from the membrane. In a final step, the Tight Helix disassembles, leading to the release of the cargo-containing vesicle.

To test the effect of cargo size on scission, we repeat the same simulation with different cargo sizes $r_{\text{cargo}} = \{1\sigma, 4\sigma, 6\sigma, 10\sigma\}$. The associated membrane-cargo interaction strength are $\{12k_{\text{B}}T$, $1.8k_{\text{B}}T$, $0.8k_{\text{B}}T$, $0.5k_{\text{B}}T\}$, respectively.

To test the effect of membrane bending rigidity on the membrane remodelling events, we repeat the same simulation with $\mu = 5$ in the membrane model, which yields a more rigid membrane with rigidity $\sim 40k_{\text{B}}T$.

When the cargo is absent, the same simulation protocol as described in the section above is adopted, except that we do not include the cargo particle.

##### S2.3 Simulation of co-equilibration of the Spiral and Wide Helix

To prepare the system, both filaments are initialized as loosely-packed spirals in the flat state, with each filament constructed of 186 subunits. A layer of 7360 membrane beads ($80\times80\sigma^2$) are positioned under the filaments in the x-y plane. Note that this initial configuration and part of the simulation described in this section are reused from a previous study [1], where the Helix is the outer strand, yet this should not change our conclusions on copolymer rigidity and stability. Initially, the membrane is equilibrated for $10^4$ steps while the protein filaments are kept frozen. The protein filaments are then released with both filaments in the flat state for about $10^6$ steps.

For the study of filament rigidity (Fig. 2 A), target radii for the Spiral and the Helix are kept constant at $R_1 = 13.2$ nm, $R_2 = 15.8$ nm, respectively, while we vary the bond strengths of the two filaments and run each simulation for around $10^6$ steps. Typically, a stable deformation is formed within $10^5$ steps. Finally, the Spiral is disassembled and only the Helix is kept on the membrane for another $10^6$ steps.

To test the stability of the copolymer scaffold (Fig. 2 C), we keep the target radius of both strands at $R_1 = 19.4$ nm initially and instantaneously change the target geometry of the Helix to the tilted state at different target radii $R_2$ for different simulations. After around $10^6$ steps of each simulation, we check whether both filaments remain attached to the membrane.

#### S2.4 Simulation of co-equilibration of the Wide Helix and Tight Helix

To test the stability of the copolymer scaffold (Fig. S3), we restart from one snapshot from the cargo-containing simulations, when the Wide Helix and the Tight Helix form a copolymer. The target radii of both helices are initially set to different $R_2$ and equilibrated for $10^6$ steps. Then the target radius of the Tight Helix is instantaneously decreased to $R_3 = 5.3$ nm and we let the system equilibrate for $10^5$ steps, before checking whether both filaments remain attached to the membrane.

#### S2.5 Simulation of the Tight Helix constriction and disassembly

To prepare the system, we place a single ESCRT-III filament on the membrane (16560 membrane beads, $120 \times 120\sigma^2$) with the cargo particle in its middle, and carry out the simulation with the filament in the flat state for $2 \times 10^4$ steps. We then switch the filament to a tilted state of target radius $R_{\text{init}} = 17$ nm and run the simulation for $\sim 2 \times 10^6$ steps. We simulate the filament constriction by progressively reducing its target radius from $R_{\text{init}}$ to $R_{\text{final}}$ at a constant rate of constriction $r_{\text{constriction}}$. The rate of constriction is defined as $r_{\text{constriction}} = (R_3[t] - R_3[t + 10\tau])/10\tau$ for any time $t$, where $R_3$ is the target radius of the Tight Helix, $\tau$ is the MD time unit. The filament is disassembled after constriction and four protocols for disassembly are tried (i.e., "instantaneous", "random", "from top", "from bottom").

The scission appears to be robust against cargo leakage. As a sanity check, we substitute the single generic cargo (radius $r = 8\sigma$) with six small cargos ($r = 2\sigma$), once the neck is thin enough to be able to sterically confine the cargos inside the budding vesicle. Volume-exclusion interactions are applied between the cargo particles (interaction strength $\epsilon = 2k_{\text{B}}T$) and short-ranged L-J interactions are applied between the cargo particles and the membrane beads (interaction strength $\epsilon = 5k_{\text{B}}T$). We carry out five replica simulations and find that four of them are successful in scission (see typical trajectory in Fig. S6). In the unsuccessful scission trajectory, the membrane retracts after disassembly of the Tight Helix filament and the cargos are not leaked out either.

#### S2.6 Mapping time unit from MD unit ($\tau$) to physical unit ($\mu s$)

The mapping expression between physical and simulation units of time is obtained by comparing the diffusion constant measured in our computer model to experimental values. To this end, we take a layer of 29440 membrane beads ($160 \times 160\sigma^2$) and equilibrate the system for $10^4$ steps with the same MD setup as detailed in the previous paragraphs. We then run the equilibration trajectory for another $3 \times 10^5$ steps and measure the mean squared displacement (MSD) of a membrane bead in the 2D membrane layer. Using the relation D= MSD/4$t$, we obtain the in-plane diffusion constant of the membrane bead D= $0.06\sigma^2/\tau$. Mapping our computed diffusion constant to the typical value for the diffusion constant of a lipid molecule in a bilayer membrane ($20\mu m^2/s$ [8]) gives our MD time unit $\tau \sim 0.018\mu s$. A similar timescale is obtained when mapping the diffusion constant of a freely-moving three-beaded protein unit in 3D (D= MSD/6$t$ = $0.25\sigma^2/\tau$) to the diffusion constant of small soluble proteins (D=$100\mu m^2/s$ [9]), which gives $\tau \sim 0.013\mu s$. However, note that one bead in our model represents a patch of lipids rather than a single lipid molecule, and our three-beaded protein unit does not necessarily represent a single ESCRT protein but rather a segment of the ESCRT polymer. Therefore, the timescale obtained above ($\tau = 0.02 \mu s$) only serves as a lower bound.

### S3 Analysis Protocols

#### S3.1 Characterization of buckle deformation

The membrane deformation is characterized by the semi-cone angle $\theta$, which is defined as the angle between the vertical axis (normal to the original flat membrane plane) and the membrane normal at half depth of the deformation. For the one-bead-thick membrane model, we take the dipole directions of the beads as the norm directions. The snapshots are saved every 1000 steps and the last 200 snapshots of the simulation are used to calculate the thermodynamic average for each simulation. The numerical values are averaged over five simulations with independent velocity seeds.

#### S3.2 Measurement of neck radius

Throughout this work, the neck radius is defined as the distance from the middle of the neck, to the inner surface of the membrane beads. To obtain it, we first measure the membrane neck radius from the centre of the neck to centre of the membrane beads. The snapshots are saved every 10,000 steps along the constriction trajectories. For each snapshot, we first extract the centre of mass of the Tight Helix in the vertical direction, $z_{\mathrm{TH}}$. The neck is then sliced into rings of thickness $\sigma$ and the diameter of each ring is estimated by the maximal distance between any two membrane beads within this ring. The neck radius is then estimated by averaging over half of the diameter of these rings that are located within $z_{\mathrm{TH}} \pm 2\sigma$. Finally, $0.5\sigma$ is deducted from the raw neck radius to obtain the inner surface membrane neck radius.

#### S3.3 Calculation of membrane potential energy profile

The bending and stretching energy between each pair of membrane beads, the membrane-filament adhesion energy between a membrane bead and a filament bottom bead, and the membrane-cargo adhesion energy between a membrane bead and the cargo bead are computed between each pair of beads in the system and evenly distributed between these two beads. The total potential energy of each membrane bead is obtained by summing up all of its pair interaction energies with all other beads. The three decomposed membrane potential energy profiles (i.e., bending and stretching, membrane-filament, membrane-cargo) are obtained by summing up only specific type of pair-pair interactions between each membrane bead and all other beads. The potential energy profiles are then binned along the membrane neck, using the bin size $\sigma$ and averaged over eight to ten snapshots after binning. Finally, the total potential energies shown in Fig. 3C are applied a constant shift so that the flat membrane has energy around zero. For visualization purposes, surface mesh is constructed by Ovito [10] using Gaussian density method [11] with resolution = 400 and iso value = 0.7.

#### S3.4 Quantification of scission efficiency

Membrane scission is detected by a DBSCAN clustering algorithm (eps=1.4, min_samples=1) since the membrane undergoes a topological transition from one cluster to two during the scission event. To quantify the scission efficiency, we carry out 50 to 60 independent constriction and disassembly simulations. Ten independent simulations are used to define a measurement and the scission efficiency is calculated by counting the number of successful scission trajectories out of those ten. We then obtain the scission efficiency by averaging over five to six independent measurements.

Table S1: Summary of all bead-bead interactions (A-B) and their default interaction strength.

| **Type of bead A** | **Type of bead B** | **Interaction Type** | **Interaction strength** |
|---|---|---|---|
| Membrane beads | membrane beads | Yuan et al.* | $\epsilon = 4.34k_{\mathrm{B}}T$ |
| Protein beads from adjacent subunits in the same filament | | harmonic bonds | $k = 256k_{\mathrm{B}}T/\sigma^2$ |
| Beads within the same three-bead protein subunit | | rigid body | NA |
| Top beads in the three-bead protein subunit | protein beads | volume exclusion | $\epsilon = 2.0k_{\mathrm{B}}T$ |
| Top beads in the three-bead protein subunit | membrane beads | volume exclusion | $\epsilon = 2.0k_{\mathrm{B}}T$ |
| Bottom beads in the three-bead protein subunit | membrane beads | Lennard Jones | $\epsilon = 3.0k_{\mathrm{B}}T$ |
| Protein bottom beads at the interfaces of different filaments | | Lennard Jones | $\epsilon = 3.0k_{\mathrm{B}}T$ |
| Protein bottom beads that are not at the filament interfaces | | volume exclusion | $\epsilon = 2.0k_{\mathrm{B}}T$ |
| Cargo bead | protein beads | volume exclusion | $\epsilon = 2.0k_{\mathrm{B}}T$ |
| Cargo bead | membrane beads | Lennard Jones | $\epsilon = 0.6k_{\mathrm{B}}T$ |

* Ref. 38 in the manuscript.

Table S2: Conversion table to map values from MD units to physical units.

| quantity | MD unit | physical unit |
|---|---|---|
| length | $\sigma$ | 2.3 nm [a] |
| time | $\tau$ | 0.02 $\mu s$ |
| energy | $k_{\mathrm{B}}T$ | 0.6 kcal/mol [b] |

[a] Taken from Harker-Kirschneck et al. (Ref. 24 in the manuscript).

[b] Assuming $T$=300K.

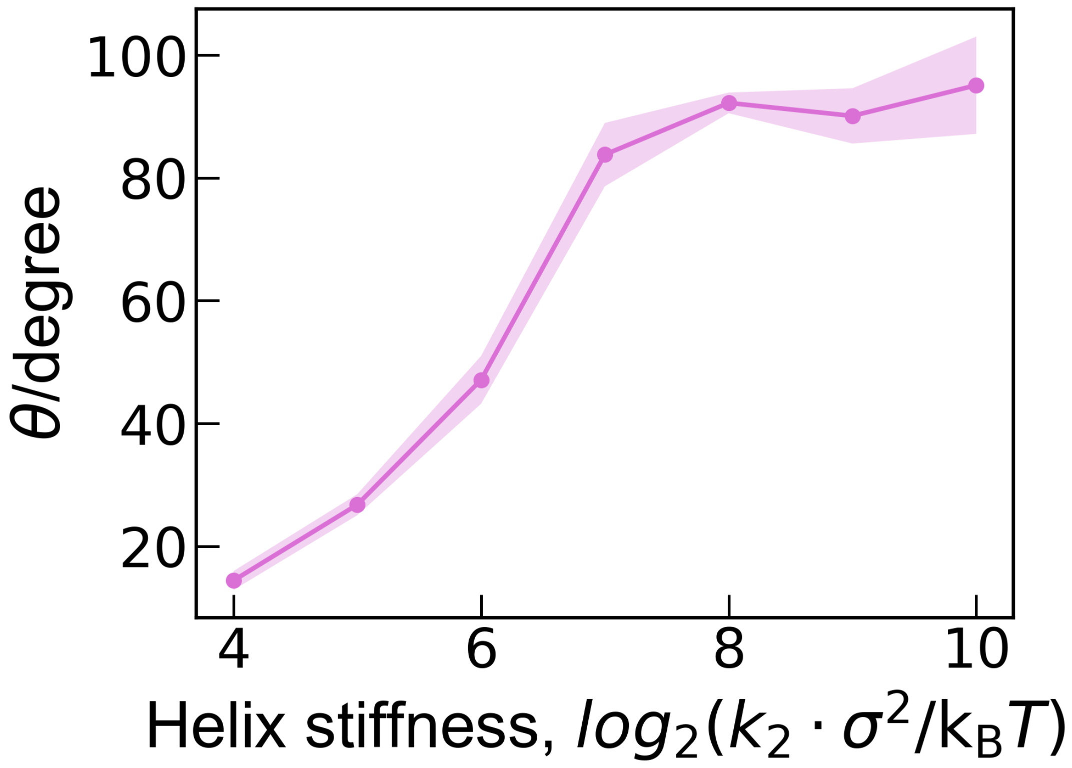


Figure S1: Membrane deformation as a function of bond stiffness of the Wide Helix ($k_2$). The membrane deformation angle $\theta$ is averaged over 5 independent simulations with the standard deviation indicated by the shaded area.

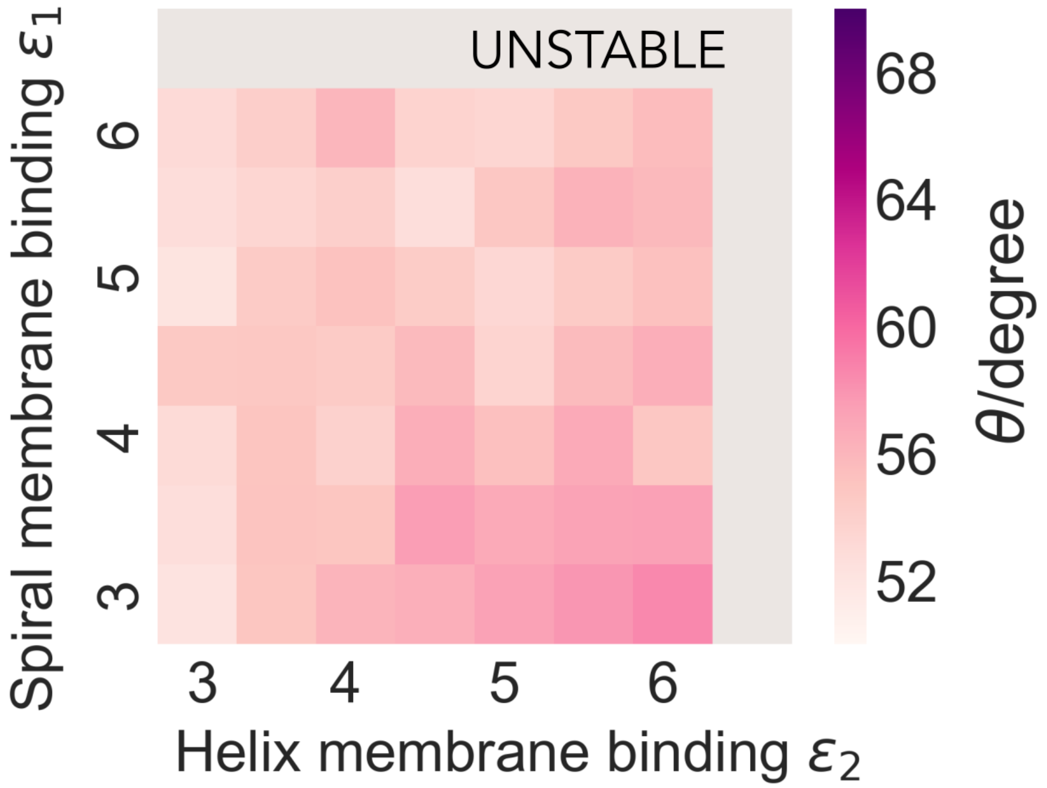


Figure S2: Membrane deformation as a function of membrane-binding affinity of the Spiral ($\epsilon_1$) and the Helix ($\epsilon_2$), ($\epsilon_1, \epsilon_2$ are in the units of $\mathrm{k_B}T$). Filament stiffness is set as $k_1 = k_2 = 256\ \sigma^2/\mathrm{k_B}T$. The membrane deformation is characterized by the angle between the vertical and the membrane norm at half depth of the deformation, as illustrated in Fig 2B. The membrane deformation is averaged over 5 independent simulations.

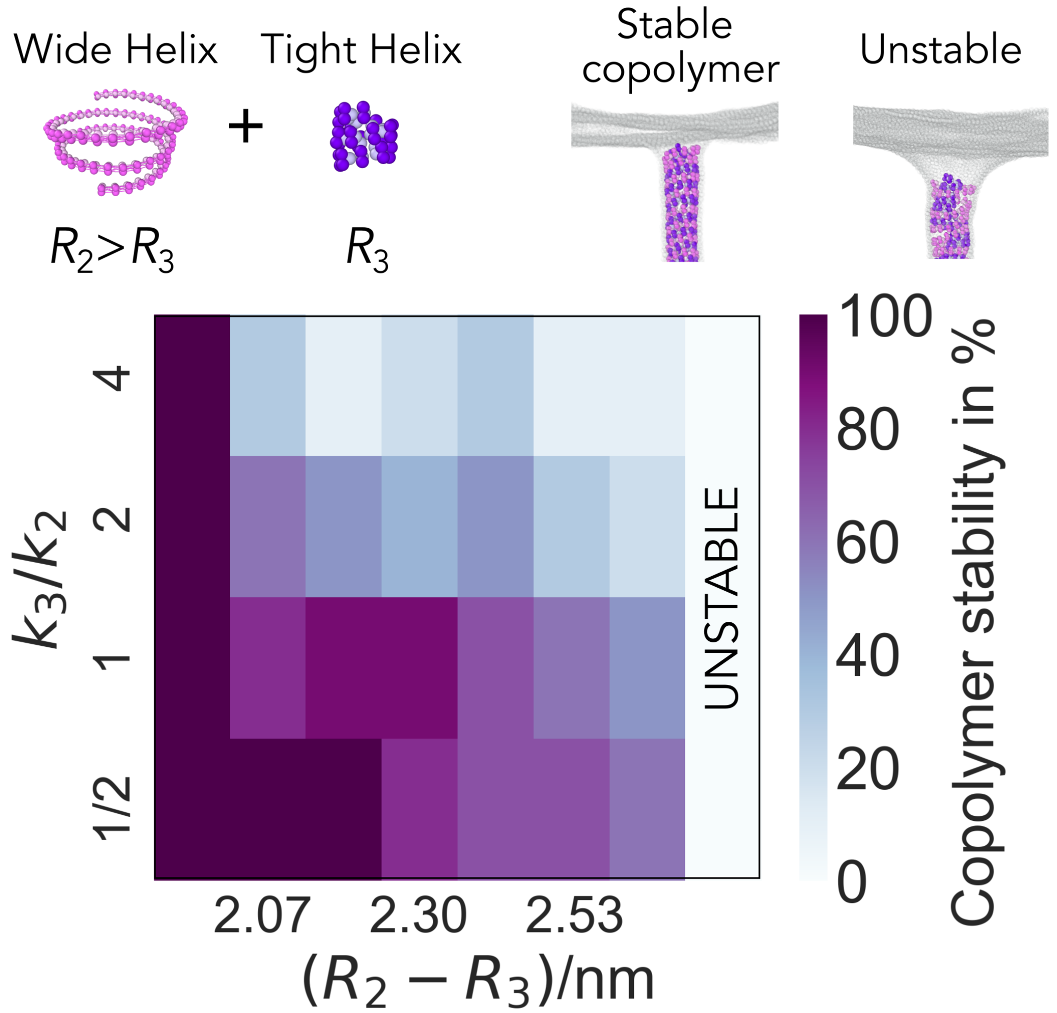


Figure S3: Copolymer stability as a function of the mismatch in the radius and ratio of stiffness between the Tight Helix and the Wide Helix. All data points are averaged over ten simulations. The Tight Helix is fixed at radius $R_3 = 5.3$ nm with filament bond stiffness $k_3 = 256\ \mathrm{k_B}T/\sigma^2$.

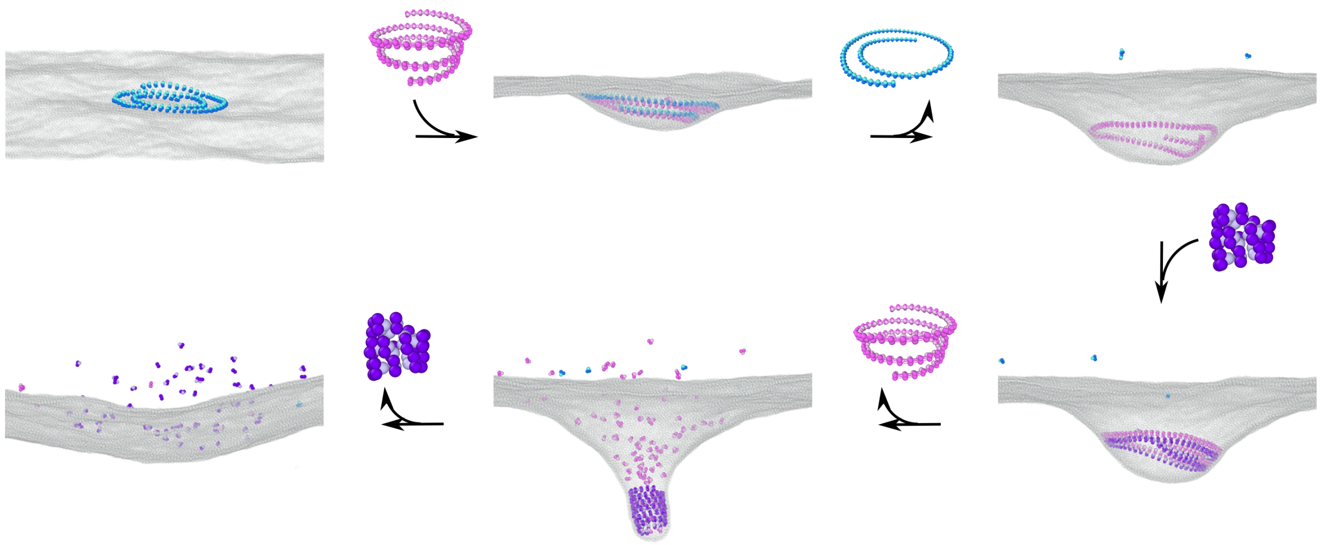

Figure S4: Typical snapshots along the trajectory where the three filaments are activated and disassembled in a stepwise manner in the absence of cargo.

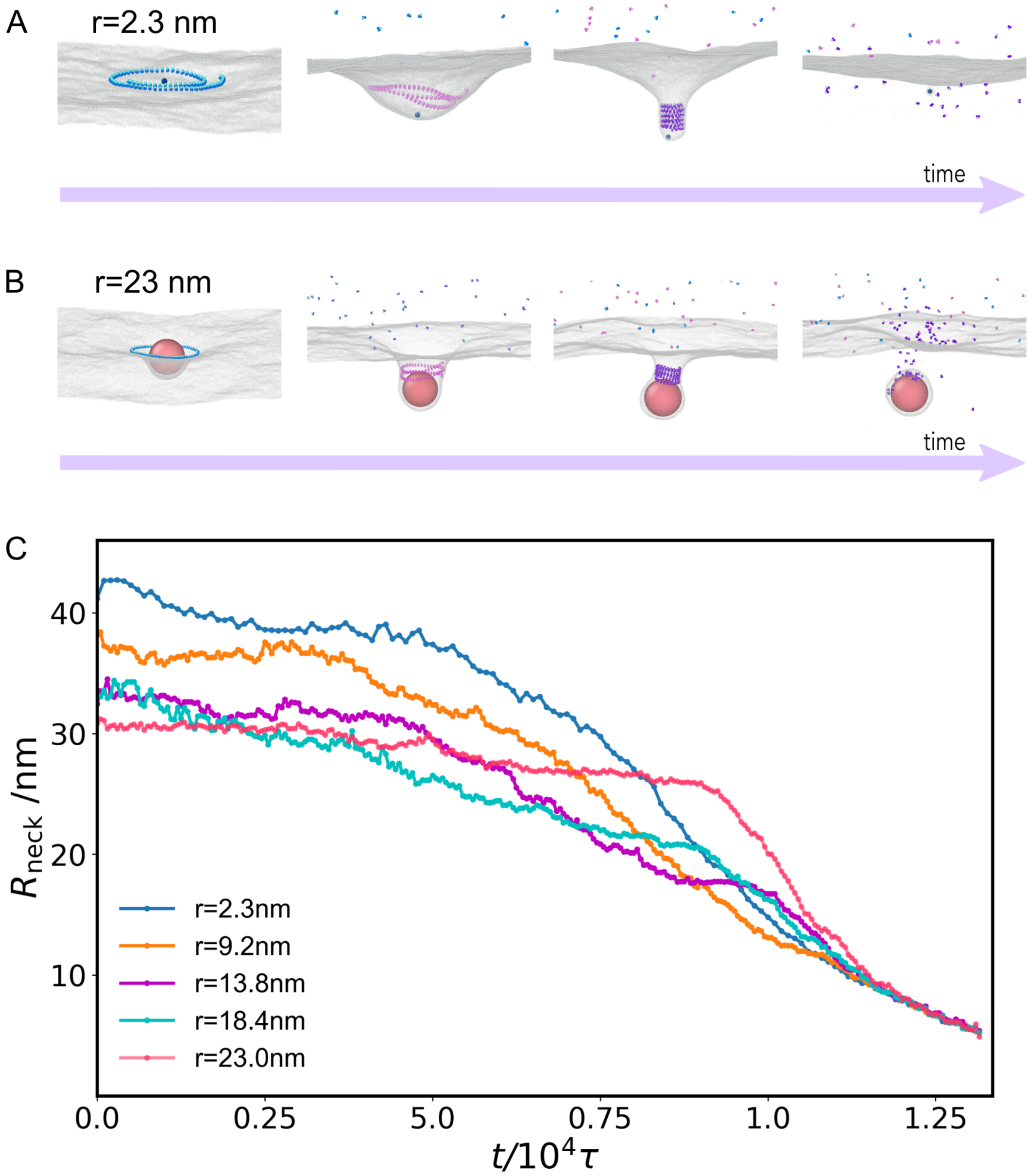


Figure S5: Impact of cargo size on the sequential recruitment and disassembly of different ESCRT-III filaments. Panels A, B: Snapshots along the trajectories of the system where the three filaments are activated and disassembled in a stepwise manner, with the radius of the generic cargo set as 2.3 nm (A) and 23 nm (B), respectively. C: Neck radii along the constriction trajectories for different cargo sizes.

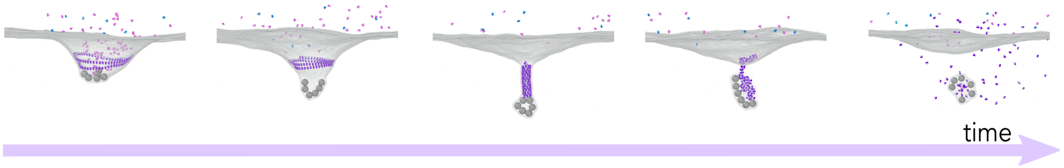


Figure S6: Tight Helix constriction and scission with six smaller volume-excluded cargos, rather than one single large volume-excluded cargo. Cargo radius $r_{\mathrm{cargo}} = 2\sigma$.

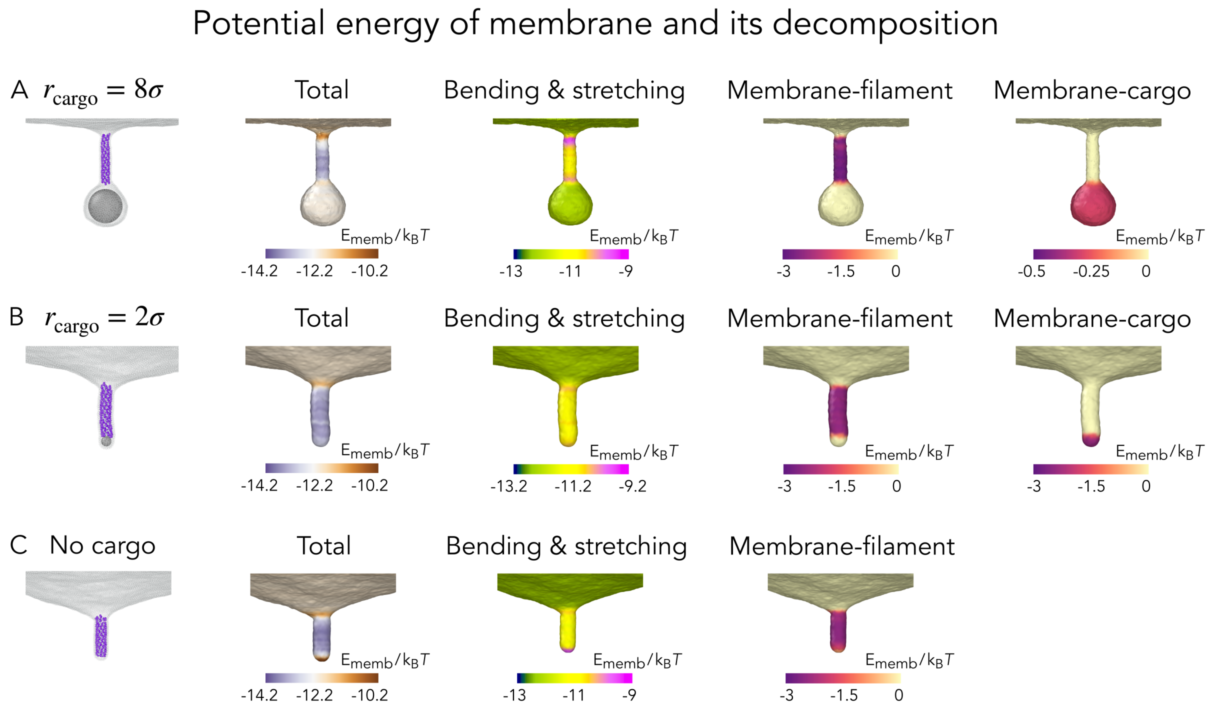


Figure S7: The local pair potential energy (i.e., energy per bead) of the membrane computed on representative snapshots before membrane breakage in the presence of the generic cargo with radii $r_{\text{cargo}} = 8\sigma$ (A), $r_{\text{cargo}} = 2\sigma$ (B), and in the absence of the generic cargo (C). The total potential is the sum of the following three potential energy terms, which are plotted separately: bending and stretching mechanical energies computed from Yuan et al. (Ref. 38 in the manuscript), filament-membrane and cargo-membrane adhesion energies computed from short-ranged LJ potential. The energies are binned along the neck of the tube with bin width $\sigma$ and averaged over 10 snapshots.

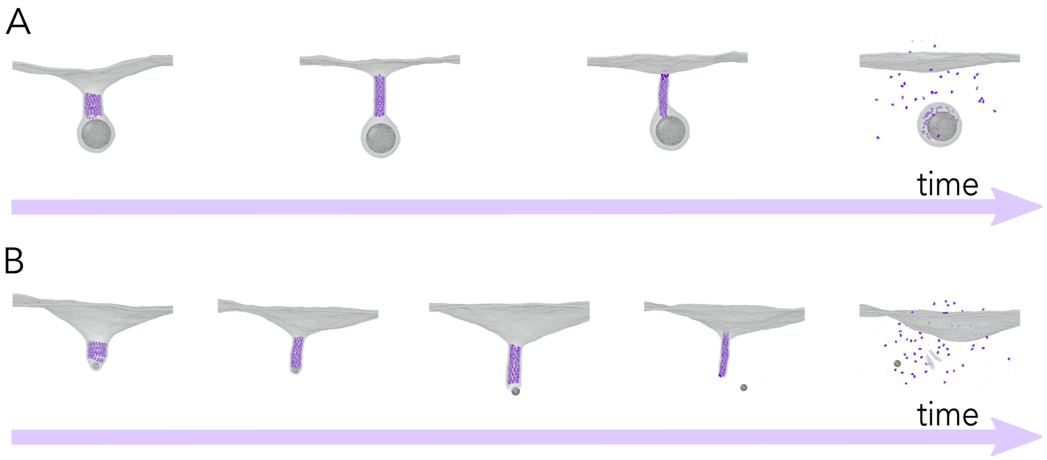


Figure S8: Tight Helix constriction and scission when the cargo-membrane adhesion is replaced by pure volume-exclusion. This change in interaction is applied once the membrane neck is thin enough to be able to sterically confine the cargo particle inside the budding vesicle or at the tip of the invagination. A: Cargo size $r_{\text{cargo}} = 8\sigma$. Successful scission is achieved with membrane breakage at the top rim of the neck. B: Cargo size $r_{\text{cargo}} = 2\sigma$. Membrane breaks at the bottom tip of the membrane neck, resulting in cargo leakage and membrane retraction.

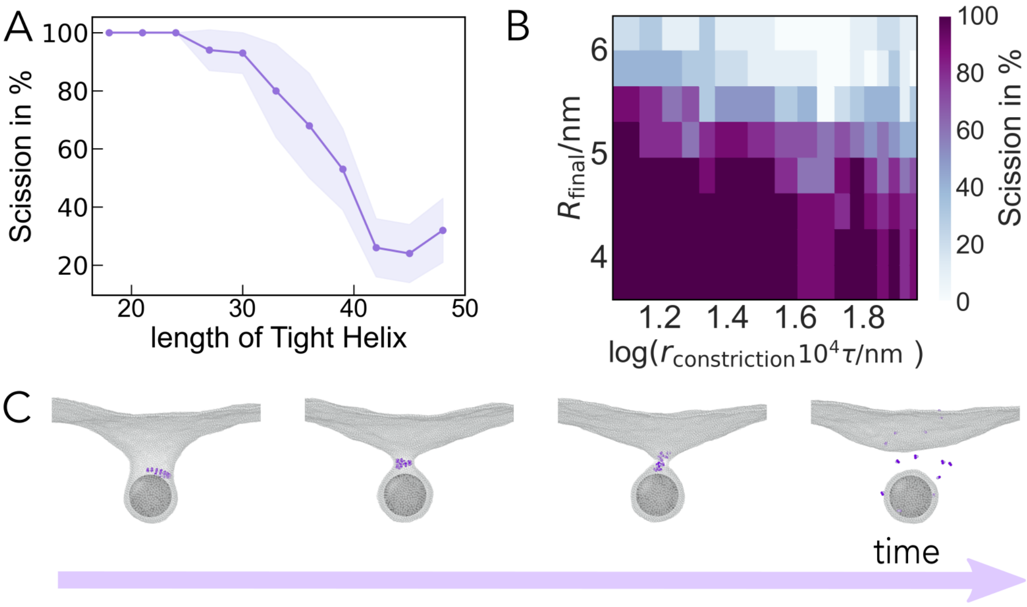


Figure S9: Shortening the length of the Tight Helix promotes scission. A: Scission efficiency as a function of the length of the Tight Helix (in number of monomers) following the protocol of progressive Tight Helix constriction followed by instantaneous disassembly. All the data is collected for $r_{\text{constriction}} = 2.1 \times 10^{-3}\text{nm}/\tau$ and $R_{\text{final}} = 4.3$ nm. The data is averaged over 5 independent measurements with the standard deviation indicated by the shaded area. B: Scission efficiency as a function of the final target radius of the Tight Helix containing 13 monomers, $R_{\text{final}}$, and the rate of the Tight Helix constriction, $r_{\text{constriction}}$. Each data point is computed from 10 independent simulations. C: Representative snapshots along the scission trajectory of the 13-monomer Tight Helix. $R_{\text{final}}$=5.6 nm and $r_{\text{constriction}} = 9.2 \times 10^{-3}\text{nm}/\tau$.

#### List of videos

- **Video 1** Stepwise activation and disassembly of the three filaments in a stepwise manner in the presence of the cargo.
- **Video 2** Stepwise activation and disassembly of the three filaments in a stepwise manner in the presence of the cargo using membrane rigidity $\kappa \sim 40 k_{\mathrm{B}}T$ ($\mu = 5$ in the membrane model).
- **Video 3** Stepwise activation and disassembly of the three filaments in a stepwise manner in the absence of the cargo.
- **Video 4** Small pores form and reseal multiple times before scission. The video pauses for a second when pores are detected.
- **Video 5** Fast Tight helix constriction: most trajectories fail in the fission. Rate of constriction $r_{\mathrm{constriction}} = 6.9 \times 10^{-3}\mathrm{nm}/\tau$ is used.
- **Video 6** Slow Tight helix constriction: most trajectories achieve fission. Rate of constriction $r_{\mathrm{constriction}} = 2.1 \times 10^{-3}\mathrm{nm}/\tau$ is used.

#### References for Supporting Information

[1] Pfitzner AK, Mercier V, Jiang X, Moser von Filseck J, Baum B, Šarić A, et al. An ESCRT-III Polymerization Sequence Drives Membrane Deformation and Fission. Cell. 2020;doi:10.1016/j.cell.2020.07.021.

[2] Harker-Kirschneck L, Baum B, Šarić A. Changes in ESCRT-III filament geometry drive membrane remodelling and fission in silico. BMC Biology. 2019;doi:10.1186/s12915-019-0700-2.

[3] Yuan H, Huang C, Li J, Lykotrafitis G, Zhang S. One-particle-thick, solvent-free, coarse-grained model for biological and biomimetic fluid membranes. Phys Rev E Stat Nonlin Soft Matter Phys. 2010;82(1). doi:10.1103/PhysRevE.82.011905.

[4] Henne WM, Buchkovich NJ, Zhao Y, Emr SD. The endosomal sorting complex ESCRT-II mediates the assembly and architecture of ESCRT-III helices. Cell. 2012;doi:10.1016/j.cell.2012.08.039.

[5] Shen QT, Schuh AL, Zheng Y, Quinney K, Wang L, Hanna M, et al. Structural analysis and modeling reveals new mechanisms governing ESCRT-III spiral filament assembly. J Cell Bio. 2014;doi:10.1083/jcb.201403108.

[6] Kozlovsky Y, Kozlov MM. Stalk model of membrane fusion: Solution of energy crisis. Biophys J. 2002;82(2). doi:10.1016/S0006-3495(02)75450-7.

[7] Plimpton S. Fast parallel algorithms for short-range molecular dynamics. J Comput Phys. 1995;117(1). doi:10.1006/jcph.1995.1039.

[8] Fahey PF, Webb WW. Lateral diffusion in phospholipid bilayer membranes and multilamellar liquid crystals. Biochemistry. 1978;17(15):3046. doi:doi: 10.1021/bi00608a016.

[9] Young ME, Carroad PA, Bell RL. Estimation of diffusion coefficients of proteins. Biotechnol Bioeng. 2010;22(5):947–955. doi:10.1002/bit.260220504.

[10] Stukowski A. Visualization and analysis of atomistic simulation data with OVITO-the Open Visualization Tool. Model Simul Mat Sci Eng. 2010;18(1). doi:10.1088/0965-0393/18/1/015012.

[11] Krone M, Stone J, Ertl T, Schulten K. Fast visualization of Gaussian density surfaces for molecular dynamics and particle system trajectories. EuroVis-Short Papers. 2012;doi:10.2312/PE/EuroVisShort/EuroVisShort2012/067-071.
